# Thymidine kinase-expressing yellow fever 17D reporter virus facilitates prodrug activation and bioorthogonal labelling of infected cells

**DOI:** 10.1101/2024.10.25.620260

**Authors:** Michael B. Yakass, Sander Jansen, Viktor Lemmens, Lorena Sanchez-Felipe, Johan Neyts, Nathan W. Luedtke, Osbourne Quaye, Kai Dallmeier

## Abstract

Tracking of viral replication and tissue tropism by *in vivo* imaging can help to unveil how live-attenuated vaccines such as the yellow fever 17D (17D) work, and likewise, to understand how adverse effects develop. Here we validate 17D-TK, a reporter virus derived from 17D that expresses herpes virus thymidine kinase (TK) that specifically converts nucleoside analogues such as Ganciclovir (GCV) to induce cell death, or difluoro-EdU (dF-EdU) for bioorthogonal labelling of infected cells by Click chemistry. 17D-TK induces a cytopathic effect in infected cell cultures, as well as mortality in intracranially inoculated mouse pups in a GCV dependent manner. Preferential phosphorylation of difluoro-EdU (dF-EdU) in 17D-TK infected cells allows to selectively stain cells that support 17D replication. Prospectively, 17D-TK can be used in combination with radiolabeled tracers for real-time detection and localization of sites of active viral replication in living animals using positron emission tomography (PET).

## INTRODUCTION

Yellow fever virus (YFV) is a (re-)emerging highly pathogenic small enveloped RNA virus that is transmitted by infected mosquitos and causes prototypic viral hemorrhagic fever (Barrett, 2018). There is no specific treatment available, yet the only intervention is immunization using vaccines based on the live-attenuated YF17D (17D) strain. Recognized for its outstanding potency, possibly inducing long-lasting polyfunctional immunity, 17D is often considered paradigm and benchmark for other vaccines and new modalities tackling unmet medical needs (Pulendran, 2009; Sanchez-Felipe et al., 2024). Despite proven efficacy and wide clinical use, with >1 billion doses deployed since its development in the 1930s, detailed knowledge about cellular targets, tissue tropism and systemic biodistribution of 17D following administration *in vivo* remains scarce, with limited insight from difficult to translate *in vitro* systems (Barba-Spaeth et al., 2005; Fernandez-Garcia et al., 2016) or indirect readouts from often complex small animal models (Douam et al., 2017; Li et al., 2022). Where and how 17D replicates in the body, viral growth kinetics in different tissues and the biodistribution of 17D remains elusive. Likewise, rare yet severe adverse events that may develop as a result of unlimited replication and systemic spread of 17D (yellow fever vaccine associated viscerotropic or neurotropic disease; YEL-AVD or YEL-AND) stay poorly understood (Barrett & Teuwen, 2009). In turn, a deeper mechanistic insight in the behavior of 17D *in vivo* may help to understand vaccination and human immunity to infection in general (Pulendran et al., 2013) with great promise for improved efficacy and safety of future vaccines.

Luciferase and fluorescence reporters for bioluminescence imaging (BLI) have been applied to track the tissue tropism of viruses in studies of pathogenesis, virus countermeasures and vaccination (Li et al., 2017; Sharma & Dallmeier, 2022; Zhang et al., 2019). Though facilitating some longitudinal analysis, spatial resolution is limited, leaving these systems fraught with the challenge of a one-time point view by sacrificing study animals and isolating organs. Inevitable contamination of thus harvested tissues with viremic blood could further obscure actual niches of viral replication (Ricklin et al., 2016). Additionally, BLI reporters are impractical for use in large animals because of limited depth capacity (<1 cm) of optical imaging systems (Gordon et al., 2019). By contrast, positron emission tomography (PET) utilizes radionuclides to provide 3-dimensional (3D) spatiotemporal imaging of biochemical and molecular events to limitless depth (nano – to picomolar sensitivity) in living cells and whole animals without the need for invasive organ retrieval (Alauddin, 2018; Gordon et al., 2019; James & Gambhir, 2012). PET can be used indirectly to visualize enhanced uptake of radiolabeled fluorodeoxyglucose ([^18^F]-FDG) due to local hypermetabolism as a proxy for viral replication and inflammation (Lau et al., 2023). By contrast, direct molecular imaging of viral infections by PET has hardly been explored. Latter setup is limited to viruses that naturally encode a marker gene (Buursma et al., 2005; Harbin et al., 1979).or recombinant viruses expressing such a marker as transgene (Chefer et al., 2018). in combination with a suitable radioactive marker substrate. Thereby, the marker substrate/probe is metabolized exclusively in infected cell, and the accumulation and distribution of the thus generated metabolite detected by gamma counter, PET, or single photon emission computed tomography (SPECT) (James & Gambhir, 2012). The human dopamine-2 receptor (hD_2_R; 443 aa) and human sodium iodide symporter (hNIS; 643 aa) which are relatively large transmembrane proteins can function as transgene markers, however, for some applications herpes simplex virus thymidine kinase (HSV-TK) may be preferred, because it is a relatively small soluble protein (about 376 aa) for which corresponding substrates/probes have been well characterized (Chitneni et al., 2007).

Thymidine kinase (TK) is an essential enzyme involved in the salvage pathway of DNA synthesis; responsible for the phosphorylation of deoxythymidine to thymidine monophosphate which subsequently as thymidine triphosphate serves as substrate for incorporation into elongating DNA. This function of TK has been applied as antiviral therapy to treat herpesvirus infections, as the low-fidelity virus-encoded TK also phosphorylates nucleoside analogues such as ganciclovir (GCV) that, when incorporated in the nascent virus genomes, abrogate virus replication. The TK/GCV system has subsequently been used as suicide gene therapy, whereby dividing cells expressing HSV-TK incorporate GCV and undergo apoptosis (Tomicic et al., 2002). The sensitivity of the TK/GCV system has since been improved by using mutant equine herpes virus TK (EHV4-TK S144H) that binds and phosphorylates GCV with higher affinity and efficacy (McSorley et al., 2014).

TK has further been applied in the so-called bioorthogonal chemical reporter labelling technique to track biomolecules in their native functional state in living systems (Prescher & Bertozzi, 2005). A functional group (e.g. azide or alkyne) is incorporated into a biomolecule (DNA, RNA, protein) in its endogenous functional state, and by Click chemistry, a cycloaddition labelled probe is used to selectively track the biomolecule (Neef et al., 2015). For example, cells in the S-phase of the cell cycle that are undergoing DNA synthesis can be selectively labelled by the incorporation of 5-ethynyl-2’-deoxyuridine (EdU) (Cavanagh et al., 2011). Pathogen-specific bioorthogonal labelling has been explored (Wang et al., 2013), whereby nucleoside analogue 2’-deoxy-2’,2’-difluoro-5-ethynyluridine (dF-EdU), a variant of EdU is metabolized and incorporated in the cellular DNA of HSV1 infected cells that thus express viral encoded TK. By this means, infected cells can be differentially stained by Click chemistry whereas uninfected cells remain unstained as dF-EdU cannot be activated by endogenous mammalian TK (Neef et al., 2015).

Live-attenuated 17D has been used as viral vector to develop vaccines against a number of pathogens (Sanchez-Felipe et al., 2024), including Japanese encephalitis and dengue (Guy et al., 2010) as well as vaccine candidates to protect from Zika congenital syndrome (Kum et al., 2018), SARS-CoV-2 (Sanchez-Felipe et al., 2021; Sharma et al., 2022) and Ebola (Lemmens et al., 2023). Traditionally, preclinical development and safety assessment of such vaccine (candidates) traditionally considered use of larger animal models (such as macaques) (Monath et al., 2015). However, biodistribution studies remain difficult in such step-up models as repeated sampling is impeded by the need to sacrifice animals for organ retrieval.

In the present study we developed 17D-TK, a 17D-derived reporter virus expressing EHV4-TK S144H. Its stability and the functionality of the TK marker is demonstrated *in vitro* and in a small animal model as proof of concept using three complementary approaches: (*i*) by GCV dependent cytotoxicity in tissue culture, (*ii*) by bioorthogonal dF-EdU labelling of infected cells, and (*iii*) by enhanced mortality due to GCV in 17D-TK infected suckling mice. Together, this validates the use of TK-expressing 17D for PET imaging of small and/or large animals in pathogenesis and vaccine studies.

## RESULTS

### Construction and characterization of 17D-TK

YF17D vaccine virus (17D) was modified to express a TK reporter (17D-TK) as translational fusion to the viral polyprotein upstream of the viral capsid protein following a previously established design (Boudewijns et al., 2021; Kum et al., 2018; Sharma et al., 2020) (**Figure 1A**). The *tk* sequence used was derived from EHV4 TK S144H mutant shown to be particularly efficient and selective in phosphorylating GCV for increased GCV-induced cytotoxicity, as compared to the wild-type enzyme (McSorley et al., 2014). To maintain essential RNA elements involved in viral genome synthesis, 63 nucleotides (corresponding to codons 1-21 of the 17D capsid gene) were inserted upstream of the *tk* gene; a C-terminal ribosome skipping 2A peptide (coT2A) was added to separate nascent TK from the rest of the 17D proteins.

**FIGURE 1.**
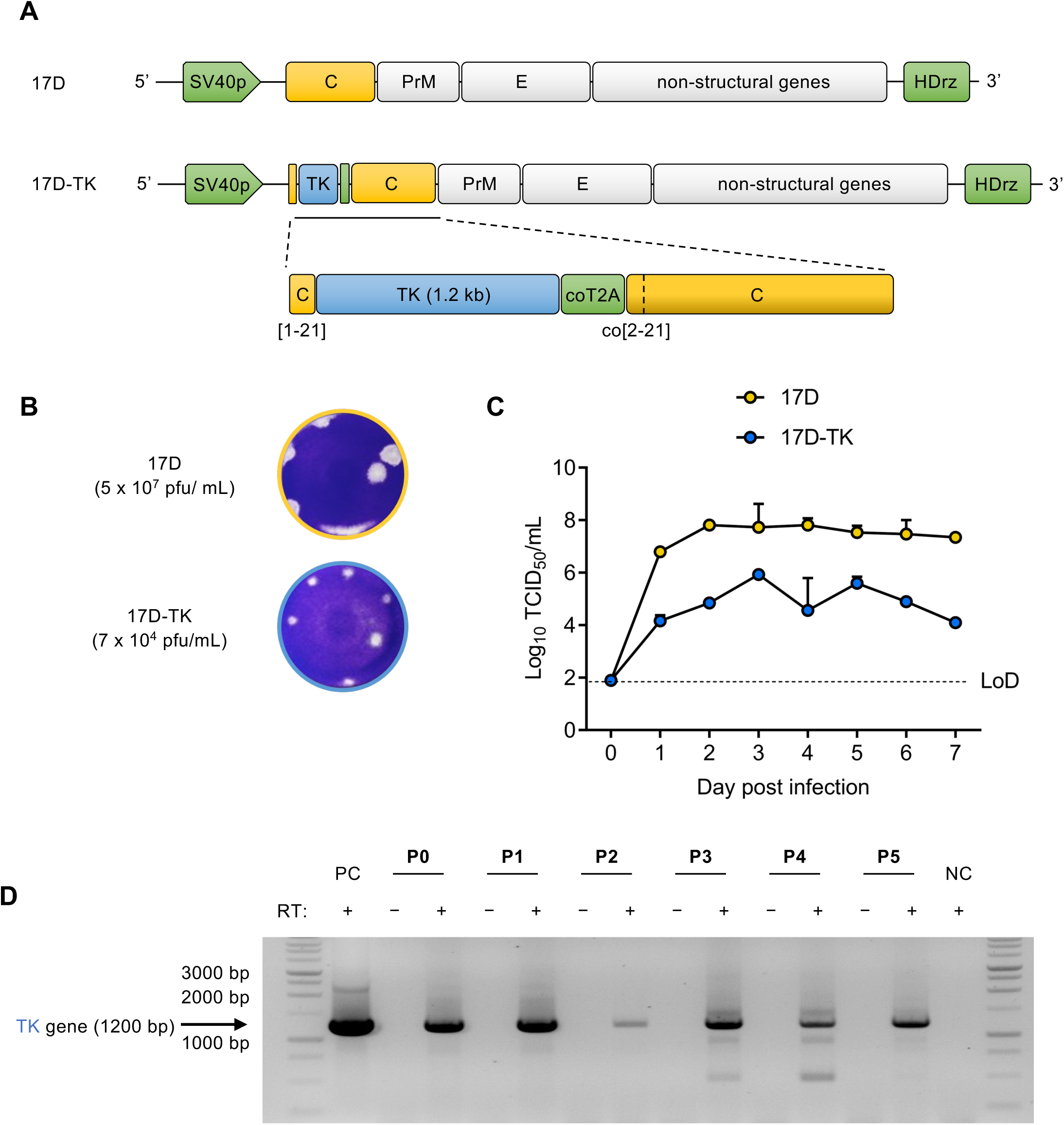
Characteristics of 17D-TK virus compared to parental 17D. **(A)** 17D-TK virus is launched from an infectious cDNA clone with the thymidine kinase (TK) gene of EHV4 inserted in the capsid gene of 17D. **(B)** Plaque phenotype of 17D-TK and parental 17D **(C)** 17D-TK and 17D were both infected at MOI 0.1 and growth kinetics were monitored by TCID50. **(D)** 17D-TK was serially passaged for five times (P1-P5) and the presence of the transgene was assessed by RT-PCR. PC, plasmid (specific amplicon 1200bp); NC, no template.

Recombinant viruses were rescued by plasmid transfection into BHK-21J cells, supernatant harvested at onset of a characteristic virus-induced cytopathic effect (CPE) and further characterized. Resulting 17D-TK virus was fully replication competent yet showed a smaller plaque phenotype (**Figure 1B**), slower growth kinetics and lower peak virus yields than parental 17D (**Figure 1C**). Genetic stability of the TK-expressing virus was assessed during serial passaging on BHK-21J and RT-PCR fingerprinting (**Figure 1D**). Genetic stability of the 17D-TK was confirmed by detection of the *tk* gene by RT-PCR fingerprinting for at least five passages (**Figure 1D**).

### GCV-dependent cytopathic effect in 17D-TK infected cells

Phosphorylation by TK turns GCV from an inert thymidine analog prodrug into a cytotoxic substrate for cellular DNA polymerases. 17D-TK is expected to express enzymatically active TK S144H. Hence, a more pronounced and/or more rapid development of CPE was expected in 17D-TK infected cells in the presence of GCV (**Figure 2B**). First, we determined the maximal dilution of 17D-TK and parental 17D that did not yet cause obvious CPE in 3 to 5 days post infection (dpi); whereby dilutions of more than 1/1600 did not result in any CPE (*data not shown*). For the following, we hence chose infections at a virus dilution of 1/1600 resulting in a MOI of 0.002 (**Figure 2B, C and D**), or at a 10-fold higher concentration (1/160; MOI 0.02) to demonstrate dose dependency in TK cytotoxicity (**Figure S2B, D**). When we next titrated the concentration of GCV needed to elicit cytotoxicity, a significant and stepwise reduction of viable cells was observed with increasing GCV concentration, with a more than 50% reduction in 17D-TK infected cell cultures at 25 µM GCV (p < 0.0001) (**Figure 2B** and **S1**).

**FIGURE 2.**
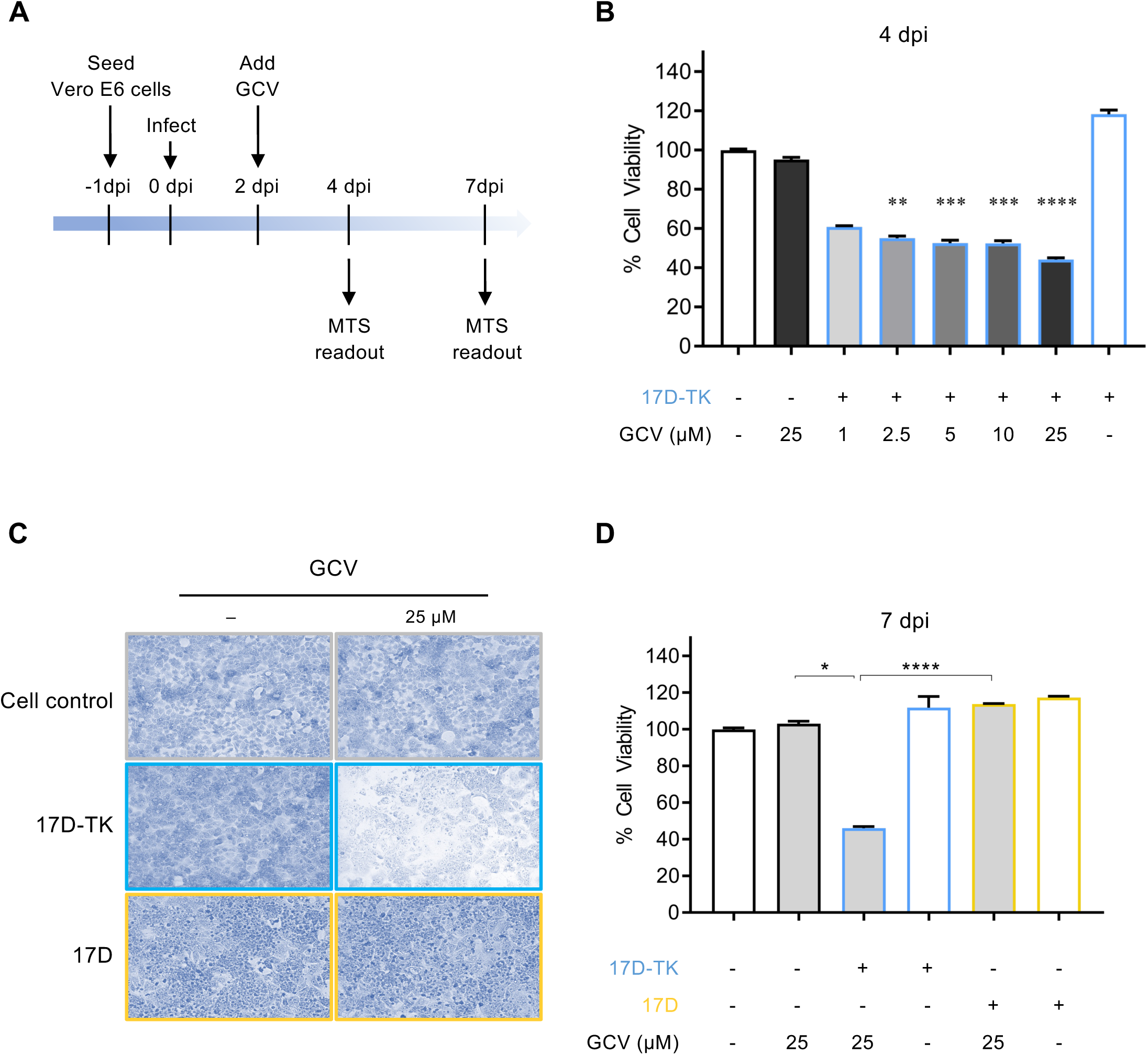
GCV induced CPE in TK-expressing cells. **(A)** Experimental set-up. **(B)** Vero E6 cells were infected with 17D-TK at a MOI of **0.002** and increasing doses of GCV applied 2 dpi. Cell viability was assessed by MTS dye conversion 4 dpi. **(C and D)** Vero E6 cells were infected **with 17D-TK or parental 17D at** MOI of **0.002**, 25 μM GCV was added 2 dpi and CPE was visualized 7 dpi by methylene blue staining of remaining cells (C; phase contrast, x40, recolored for contrast) or quantified by MTS **(D).** Uninfected cells served as controls in all cases. Bar graphs represent Mean + SEM from two independent experiments each in n = 6 technical replicates. * p<0.05; ** p<0.01; *** p<0.001; **** p<0.0001. GCV, ganciclovir; TK, thymidine kinase; dpi, days post infection.

Too early GCV treatment seemed to reduce its own effect as it may have led to early elimination of virus infected cells from the culture system (**Figure S2B**). Therefore, an optimized schedule was developed to balance TK expression and subsequent GCV treatment (**Figure S2C**). By delaying treatment for 2 dpi, the effect of GCV could be enhanced (**Figure S2D**). At 5 days post GCV application (corresponding to 7 dpi), lower doses sufficed (10 µM: Mean viability 38%, p < 0.0001), a more pronounced CPE could be achieved (25 µM: Mean viability 33%, p < 0.0001), or the effective MOI could further be reduced (MOI of 0.002: Mean viability 57%, p = 0.008). Taken together, GCV toxicity seemed to be dependent on abundant TK expression in cells transduced by the recombinant 17D-TK vector. Importantly, GCV treatment had no effect on cells infected with parental 17D as there was no unspecific GCV-induced cytotoxicity observed in 17D-infected cells even at a high GCV concentration of 25 µM nor during an extended period of 5 days (**Figure 2C** and **D**), nor a 10-fold viral titer of 17D at MOI of 0.02 (**Figure S1**).

### Effect of GCV on 17D-TK infected suckling mice

Intracranial inoculation of newborn mice with 17D is known to lead to growth retardation and mortality in a dose dependent manner. Likewise, recombinant 17D variants have been shown to have a generally attenuated neurovirulence (Lemmens et al., 2023; Monath et al., 2015; Sanchez-Felipe et al., 2024; Sanchez-Felipe et al., 2021). Indeed, when 4 – 5 days old Balb/c pups were infected with either 100 PFU of 17D and 500 PFU 17D-TK via the intracranial route (**Figure 3A**), 17D infected animals stopped growing within 3 – 4 dpi, while initial development of 17D-TK infected individuals followed that of uninfected controls; gaining on average 110% of their starting weight within 6 dpi (**Figure 3B**). By contrast, 17D infected pups had gained only 50% weight, became sick from day 6 requiring euthanasia by day 7. 17D-TK infected pups remained healthy until day 10, requiring euthanasia on day 11 (**Figure S3A**). Such a reduced neurovirulence of 17D-TK is in line with the previously observed reduced replication fitness (**Figure 1B, C**). Importantly, viral RNA loads as determined by RT-qPCR at time of euthanasia and thus reporter expression in 17D-TK animals did not vary between groups (**Figure S3F**).

**Figure 3.**
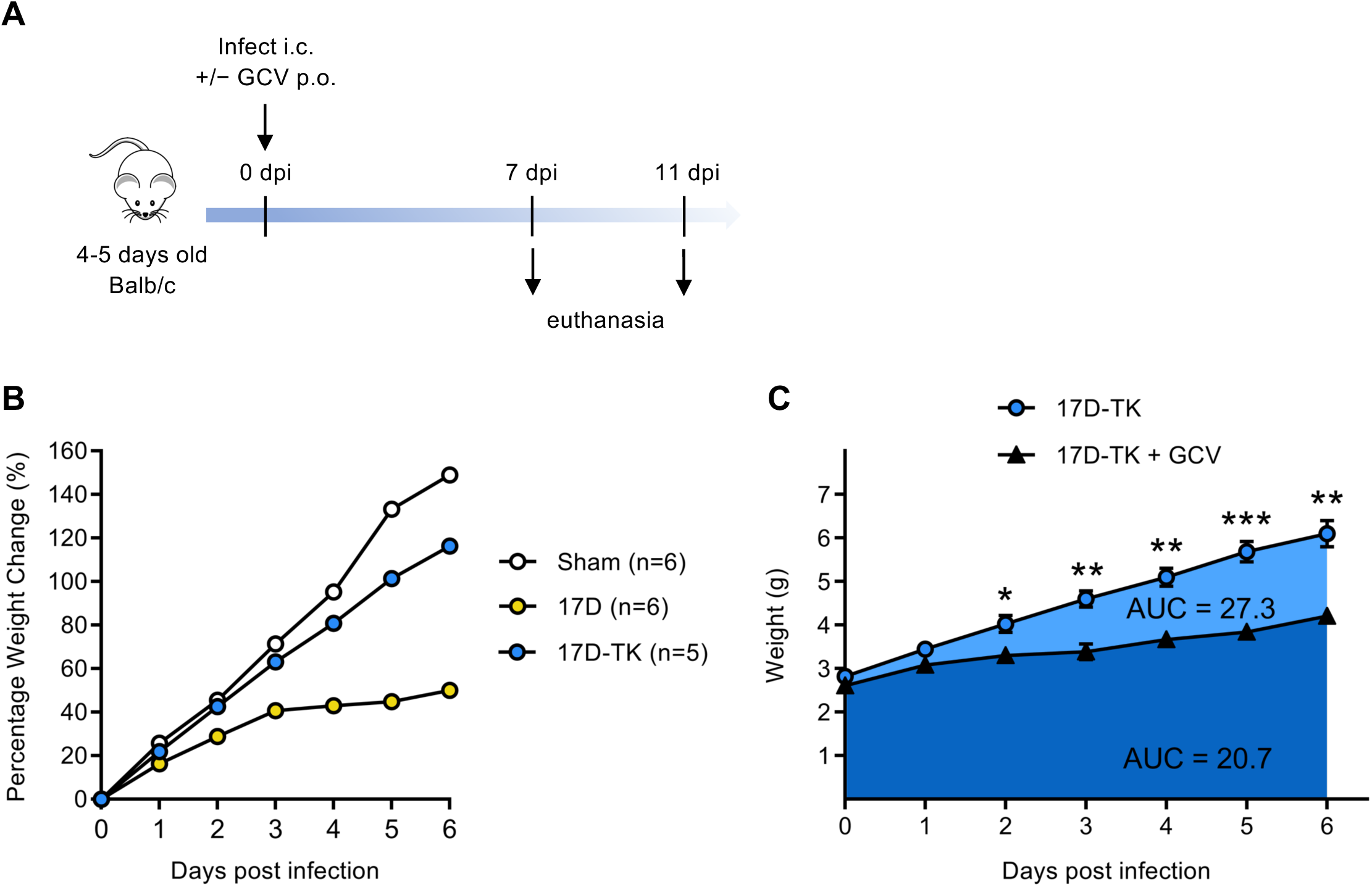
GCV dependent mortality of 17D-TK infected Balb/c pups. **(A)** Attenuated neurovirulence of 17D-TK compared to parental 17D. Balb/c pups (4 to 5 days old) were either sham infected, or infected with 100 PFU of 17D or 500 PFU of 17D-TK virus intracranially and the weight was monitored daily. **(B)** Retarded weight evolution of 17DTK infected pups when exposed to GCV *per os* (n=5) by treating foster mothers once daily with 100 mg/kg GCV intraperitoneally, compared to unexposed pups (n=5). Difference in cumulative weight calculated as mean AUC+/-SEM. Holm-Sidak t-test; * p<0.05; ** p<0.01.

This milder infection outcome and prolonged survival time in 17D-TK infected animals allowed us to assess whether GCV had a similar enhancing effect on disease progression and infection outcome *in vivo* as shown on CPE *in vitro*. GCV was dosed orally by treatment of the fostering mothers with GCV, known to be transferred via breast milk (Alcorn & McNamara, 2002) and being accessible to the brain (Biron, 2006; Field et al., 1984). Under these circumstances, GCV caused a significantly growth retardation of 17D-TK infected pups, seen a markedly flattened slope and lower AUC of their growth curves (20.7 ± 0.55; 95% CI: 19.6 – 21.7) compared to 17D-TK non-treated pups (AUC 27.3 ± 0.79; 95% CI: 25.7 – 28.9) (**Figure 3C** and **S3A, B**). Finally, over a period of 6 days, the weight of GCV treated animals (∼ 4g) reached roughly only 65% of the weight of the untreated controls (∼ 6g) (**Figure 3B**). Likewise, GCV alone was well tolerated and had no effect on uninfected pups (**Figure S3C-E**).

Overall, these data confirm the specific activation of GCV at the site of 17D-TK replication, hence, conversion of other substrates of TK should also be feasible *in vivo*.

### Bioorthogonal labelling of 17D-TK infected cells *in vitro*

EdU incorporation followed by dye-coupling via Click chemistry is a widely used method for chemical labelling of nascent DNA in cells (Salic & Mitchison, 2008). While EdU can be activated by ubiquitous cellular kinases, incorporation of its analogue dF-EdU depends on specific phosphorylation by a herpes virus TK (Neef et al., 2015). Accordingly, cells infected with 17D-TK, but neither uninfected cells nor cells infected with parental 17D used as control, exhibited nuclear staining upon the addition of dF-EdU and a fluorescent azide (**Figure 4**). Whereas general EdU labelling was achieved in dividing Vero E6 cells in cell culture, independently whether they were infected by 17D-TK, 17D or left uninfected (**Figure S4**), dF-EdU incorporation occurred exclusively in 17D-TK infected cultures; whereby dF-EdU signal frequently co-localized with a prominent staining for 17D antigens (**Figure 4A**) and dsRNA (**Figure 4B**), as markers for viral gene expression and ongoing virus replication.

**FIGURE 4.**
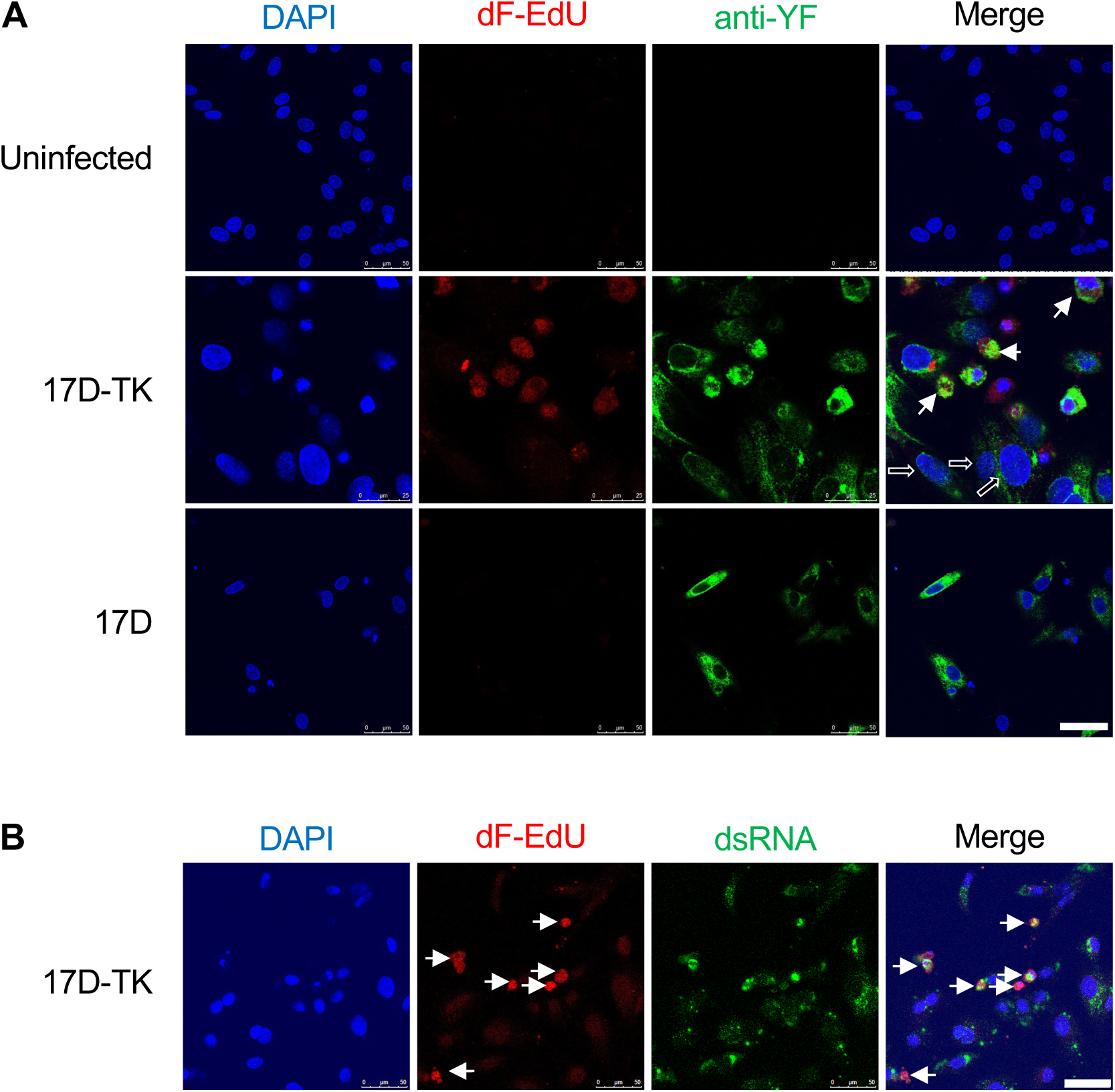
17D-TK infected cells and virus progeny incorporate dF-EdU and differentially stain by bioorthogonal labelling. Vero E6 cells were infected with 17D-TK or parental 17D at a MOI of 3 and incubated with the thymidine analogue dFEdU for two days. Cells were washed and stained by click chemistry using Alexa-fluor-594 conjugated azide. Cells were incubated with **(A)** anti-yellow fever hamster polyclonal antibody or **(B)** with anti-dsRNA and stained with Alexa Fluor-488 antimouse secondary antibody with DAPI as DNA stain. Uninfected cells served as controls. Confocal microscopy images are representative of two independent experiments. Replicating (S-phase) infected cells indicated by bold arrows in **(A)** and **(B)**, non-S phase infected cells indicated with hollowed arrows in **(A)**, co-stained virus progeny indicated with dotted circles in **(B).** I.N; indistinct nuclei, dF-EdU; 2’-deoxy-2’,2’-difluoro-5-ethynyluridine, white scale bar = 50 μm

Specific incorporation of dF-EdU by 17D-TK infected cells (**Figure 4**), combined with the observation that GCV exacerbated 17D-TK induced neurovirulence in suckling mice (**Figure 3**) suggested that 17D-TK infected tissue could be similarly identified by bioorthogonal labelling *in vivo*.

## DISCUSSION

In this report, we have validated a TK-expressing yellow fever 17D vaccine virus (17D-TK) to be used as reporter virus *in vivo*, e.g. in pathogenesis and mechanistic vaccine studies in small and large animal models. For initial characterization and first proof of concept, we tested two complementary means of detection; GCV-dependent CPE/cytotoxicity and metabolic/bioorthogonal labelling, demonstrating that 17D-TK is functional and fit for future intended studies using PET imaging with radiolabeled TK-substrates such as [^18^F]-FMAU, [^18^F]-FEAU or [^18^F]-FHPG (Alauddin, 2018; Brust et al., 2001; Chin et al., 2008).

PET imaging may enable longitudinal studies of viral infections *in vivo*, and should allow for visualization, characterization, and quantification of viruses and their dynamics in both, small and large animals (Lau et al., 2023). However, so far, most studies of viral pathogenesis using PET, including preclinical and clinical studies on HIV, dengue virus and SARS-CoV-2, employed [^18^F]-FDG uptake and cellular hypermetabolism that is associated with virus-induced local inflammation as proxy for active viral replication (Lau et al., 2023). Reports on direct visualization of infected cells by PET remain limited. One study explored recombinant MERS coronavirus expressing hNIS, but proved unsuccessful *in vivo* due to genetic instability of the reporter virus construct. In fact, the hNIS transgene was immediately eliminated from the viral RNA genome during MERS replication (Chefer et al., 2018). By contrast, our 17D-TK virus maintained its TK reporter gene for at least five serial passages *in vitro* (**Figure 1D**) as well as TK activity for at least six days of intracranial virus growth in suckling mice (**Figure 3**). Finally, cells infected with human herpes virus (HSV1 and human cytomegalovirus) have successfully been labelled using [^18^F]-FHPG as substrate for their endogenous TK. In continuation, [^18^F]-FHPG PET has been used for the detection of HSV1 in a rat model of experimental herpes encephalitis (Buursma et al., 2005; Harbin et al., 1979). Altogether, this may encourage the use of 17D-TK for PET imaging *in vivo*.

TK-dependent GCV-induced cytotoxicity has been described and applied frequently in the field of gene therapy (Beltinger et al., 2002; Chhikara et al., 2001; Sandmair et al., 2000). Hereby, herpes virus TK phosphorylates GCV to GCV-monophosphate which is further phosphorylated by host cell kinases with the resulting GCV-triphosphate (GCV-TP) incorporated into replicating DNA and leading to arrest of the cell cycle in S-phase and eventual apoptosis (Tomicic et al., 2002). As first evidence that the 17D-TK reporter virus was functional, dose-dependent cytotoxicity could be observed with up to 70% cell death observed in 17D-TK infected cell cultures when incubated with 25 µM GCV; within a reasonable concentration range previously reported for other cells (McSorley et al., 2014; Tomicic et al., 2002).

YFV is a cytopathic virus and infecting cells with 17D vaccine strain at a high MOI causes significant CPE within 2 – 3 dpi. Therefore, specific GCV-induced cytotoxicity could only be detected at a virus dose low enough not to cause any cell death until 7 dpi, yet high enough to ensure that sufficient cells were infected to express TK. After serial titrations, MOI of 0.02 and 0.002 were chosen at which, even for the more aggressively growing parental 17D (**Figure 1B**), no obvious CPE could be observed (**Figure 2, S1 and S2**).

EdU is a nucleoside analogue that can be phosphorylated by cellular TK and is widely used for bioorthogonal labelling of nascent DNA in the study of proliferation, carcinogenesis and cell cycle dynamics (Neef et al., 2012). In contrast to specific EdU incorporation by cells infected with DNA viruses such as HSV1 (Wang et al., 2013), we did not see any preferential nor differential EdU incorporation in 17D-TK or 17D infected cells (**Figure S4**). dF-EdU is a derivative of EdU that is preferentially metabolized by herpesvirus TK and differentially incorporated into nuclear DNA of cells expressing such enzymes (Neef et al., 2015). Here we demonstrate the capability of 17D-TK to activate dF-EdU for metabolic labelling exclusively of infected cells by co-staining for viral antigens and dsRNA (**Figure 4A**), validating 17D-TK as live reporter virus that allows to track viral spread in cell culture and possibly sites of virus replication in tissues.

## LIMITATIONS OF THE STUDY

There are several limitations to our study. Firstly, the enzymatic activity of TK expressed by 17D-TK is not directly measured yet only indicated by the toxicity of GCV in combination with viral infection; yet likely justified by the well characterized TK/GCV paradigm also used in suicide gene therapy. More importantly, the TK gene has been shown to be stable in 17D-TK to the fifth cell culture passage. It is unclear if this is sufficient for *in vivo* applications such as tracking of virus trajectories after peripheral inoculation from the local site of initial replication to secondary target organs that depends on massive virus growth and proliferation. Test of virus stability for long-term PET imaging as intended was outside the scope of the current study.

Finally, dF-EdU labelling of 17D-TK infected cells is demonstrated only *in vitro*. Similar *in vivo* proof may not be attainable due to inherent limitations of the infection model. While replicating virus was widely distributed in the brains of 17D-TK infected mouse pups and could readily be detected by staining for YFV dsRNA (**Figure S5**), only a small fraction of cells showed cellular DNA synthesis and, hence, incorporation of EdU, in line with strictly organized developmental patterns in the brain of newborn mice. Likewise, also dF-EdU labelling requires actively growing cells. Consequently, failure of metabolic labelling of resting cells might be misinterpreted as if 17D (17D-TK) tropism, infection and replication was selective for cells undergoing cell division. Certain areas in the brains of pups may still offer this milieu. By contrast, other relevant target cells *in vivo* such as monocytes/macrophages and dendritic cells do not divide. Latter shortcoming will only be overcome once radioactive substrates are used that, following conversion by TK, accumulate as soluble intracellular tracers within infected cells (Alauddin, 2018; Brust et al., 2001; Chin et al., 2008), independently of cell cycle phase and cellular differentiation status. Such substrates have not yet been tested as they are not abundantly available, requiring custom synthesis by specialized radiochemistry labs used to work with short-lived radioisotopes.

In conclusion, we developed a viable TK-expressing yellow fever 17D reporter virus and validated its function using two complementary approaches (suicide gene approach; bioorthogonal labelling) *in vitro* and *in vivo*. 17D-TK and similar constructs based on wild type YFV (YFV-Asibi strain) can provide powerful tools to study viral pathogenesis, vaccine attenuation and safety, and antiviral treatments using PET imaging in small and large animals.

## AUTHOR CONTRIBUTIONS

MBY and KD conceptualized the project and designed the experiments. MBY performed the experiments and wrote the initial manuscript draft. LSF constructed 17D-TK virus. MBY performed animal experiments. MBY and VL performed immunohistochemistry staining experiments. MBY and SJ performed imaging of immunohistochemistry slides and made results figures. NL provided dF-EdU. JN and KD secured funding for the research and KD supervised the project. MBY, OQ and KD finalized the manuscript with the help of all co-authors. All authors read and approved the final manuscript.

## Supporting information

Key Resources

## ACKNOWLEDGEMENTS

The authors thank Sarah Debaveye for help with molecular cloning of the 17D-TK construct and Dr. Mahadesh P.A. Javarappa for help with intracranial inoculations. This work was supported by the Flemish Research Foundation (FWO) Excellence of Science (EOS) program (No. 30981113; VirEOS project and No. 40007527; VirEOS2), and the European Union’s Horizon 2020 research and innovation program (No. 733176; RABYD-VAX consortium).

The authors thank Dr. Yeranddy A. Alpizar (MVVD) and Dr. Christopher Cawthorne, KU Leuven Department of Imaging & Pathology (Nuclear Medicine & Molecular Imaging Unit) for helpful discussions and critical comments on the manuscript.

## STAR METHODS

### RESOURCE AVAILABILITY

#### Lead Contact

Information and requests for resources and reagents should be directed to and will be fulfilled by the lead contact, Kai Dallmeier (kai.dallmeier@kul<u>euven.be</u>)

#### Materials Availability

Virus constructs made during this research may be accessed depending on availability and intellectual property arrangements by placing a request to the lead contact.

#### Data and Code Availability

Any additional information required to reanalyse the data reported in this paper is available from the lead contact upon request

### EXPERIMENTAL MODEL AND SUBJECT DETAILS

#### Animals

Balb/c pups (4-5 days old) were purchased with their mothers from Janvier Laboratories (France) and housed five to seven pups and mother in isolator cages (Biocontainment systems, Techniplast) with access to cage enrichment, food and water *ad libitum*. All animal experimental procedures were reviewed and approved by the ethical committee of KU Leuven (License: P100/2020) following institutional guidelines approved by the Federation of European Laboratory Animal Science Associations (FELASA). At both humane and experimental endpoint, pups or mothers were respectively euthanized by 20 µL or 200 µL Dolethal (200 mg/mL sodium pentobarbital, Vetoquinol SA). A group of five to seven Balb/c pups were either sham infected (media) or infected with 100 PFU of 17D or 500 PFU of 17D-TK intracranially. Weight of pups were recorded daily and monitored for morbidity or mortality.

#### Cell Lines

BHK-21J and Vero E6 cells used in this study were a generous gift from Peter Bredenbeek, LUMC, NL. Cells were maintained in seeding medium containing DMEM medium (Gibco, Belgium) supplemented with 10% foetal calf serum (FCS, Gibco, Belgium), 2 mM glutamine (Gibco, Belgium) and 0.75% sodium bicarbonate (Gibco, Belgium). Virus culture and cytopathic effect-based experiments were performed in assay medium, which is the seeding medium supplemented with only 2% FCS. All cultures were maintained at 37 °C in an atmosphere of 5% CO_2_ and 95–99% humidity.

#### Microbe Strains

17D-TK used in this study is a plasmid-launched virus constructed in our laboratory. To make 17D-TK virus, a DNA fragment encoding the equine herpes virus-4 thymidine kinase (EHV4 TK) with S144H mutation was custom synthetized (IDT Integrated DNA Technologies, Haasrode, Belgium) and subcloned into a BAC pShuttle-17D expression vector (Patent Number: WO2014174078 A1) of our Plasmid-Launched Live-Attenuated Vaccine (PLLAV)-17D platform using standard molecular biology techniques inserting EHV4 TK into the capsid (C) gene of pShuttle-17D. EHV4 TK with serine to histidine mutation at position 144 has been demonstrated to bind and phosphorylate ganciclovir much efficiently than wild type EHV4-TK and HSV1 TK (McSorley et al., 2014). To generate 17D-TK virus, BHK 21J cells were transfected with PLLAV-17D-TK construct using TransIT®-LT1 transfection reagent, following the manufacturer’s instructions. Upon onset of CPE, 17D-TK virus was harvested, centrifuged at 4000 rpm at 4 °C for 10 mins, aliquoted and stored at −80 °C. Concentration of viruses was determined by plaque assay (PFU/mL), as described below.

### METHOD DETAILS

#### Plaque assay

A day prior, 10^6^ BHK-21J cells were seeded in 6-well plates for overnight incubation. Cells were washed once with PBS, infected with ten-fold serial dilutions of virus, and incubated for 1 h at room temperature for virus attachment. Uninfected cells were used as negative controls. Cells were washed three times with the assay medium and overlaid with MEM-2X (Gibco, Belgium) supplemented with 4% FCS and 0.75% sodium bicarbonate containing 1% low melting agarose (Invitrogen, USA). When agar had set and solidified, plates were incubated for 5 days at 37 °C, 5% CO_2_. Cells and overlay were fixed with 8% paraformaldehyde, agarose was removed, and cells were stained with 1% crystal violet solution. Plaque phenotype was noted, plaques were counted, and virus concentration estimated as plaque forming unit per mL (PFU/mL).

#### Virus replication kinetics

BHK-21J cells were seeded at 2 x 10^5^ cells per well in 6-well plates for overnight incubation at 37 °C, 5% CO_2_. Cells were washed with PBS and infected with 17D-TK or parental 17D at MOI of 0.1 and incubated for 1 h at 37 °C. Cells were then washed twice with PBS and cultured with DMEM (Gibco, Belgium) supplemented with 2% FCS at 37 °C, 5% CO_2_. Culture supernatants were collected daily for six days, centrifuged at 2000 rpm for 10 mins, aliquoted and stored at −80 °C until use. Virus titer was estimated by determining the 50% tissue culture infectivity dose (TCID_50_), as described below.

#### Virus titer estimation by TCID_50_

BHK-21J cells (2 x 10^4^) were seeded in 96-well plates overnight. The following morning, seeding medium was changed to assay medium and ten-fold serial dilutions of virus samples were made in 96-well plates and cultured at 37 °C, 5% CO_2_ for 5 days. The plates were then observed microscopically for CPE and stained with MTS/phenazine methosulphate (PMS; Sigma-Aldrich) solution for 1 h at 37 °C in the dark. After this, the absorbance was measured at 498 nm for each well. Assays were performed in six-replicates and TCID_50_ per mL was determined using the Reed and Muench method (REED & MUENCH, 1938).

#### Virus induced CPE titration

We determined a virus concentration/MOI for both parental 17D and reporter virus 17D-TK that did not produce any obvious CPE by 7dpi. Vero E6 cells (10^4^) were seeded in 96-well plates and the first well infected with undiluted virus stock. Wells in the first row were then infected from this first well creating a log-2-fold serial dilution across the first row of wells. Subsequent rows of wells were infected by creating a second log-5-fold serial dilution down the plate from the first row of infected wells. Plates were incubated at 37 °C, 5% CO_2_ and wells observed for CPE daily for 7 days. Wells without obvious CPE on 7 dpi were noted and the virus titer in these wells were computed.

#### TK/GCV-enhanced cytotoxicity assay

Two different setups were made to determine the combined effects of thymidine kinase-expressing virus (17D-TK) and GCV application. In one setup, Vero E6 cells were seeded in 96-well plates at 10^4^ cells per well for overnight incubation. Cells were infected with 17D-TK or parental 17D virus at MOIs of 0.02 or 0.002 and GCV was added to wells at final concentrations of 10 µM or 25 µM. As controls, and to delineate the combined and or singular effect of virus or GCV, 10 µM or 25 µM of GCV was added to uninfected cells as compound control, while virus-infected cells served as virus control. Plates were incubated at 37 °C, 5% CO_2_ and MTS assay performed at 3 and 5 days post infection. In the second setup, similar workflow was followed, however, GCV was applied at 2 dpi and MTS performed 4 dpi (2 days post GCV application) and 7 dpi (5 days post GCV application). Two independent experiments were performed with each test condition performed in 6 replicates.

#### TK/GCV-induced CPE staining assay

Vero E6 cells (10^4^) were seeded in 96 wells for overnight incubation. Cells were infected with 17D-TK or parental 17D virus at MOI 0.02 or 0.002 with each concentration in 6 replicate wells and incubated at 37°C, 5% CO_2_. Uninfected cells served as negative controls. GCV was applied at 2 days post infection to respective wells (17D-TK, 17D or uninfected) at a final concentration of 10 µM or 25 µM. Plates were incubated at 37 °C, 5% CO_2_ for 5 more days. Cells were washed twice with PBS, fixed with 70% ethanol and stained with methylene blue. Two independent experiments were performed, and each experimental condition assayed in 6 replicates. Morphology and or cytopathic effect (CPE) of stained monolayer of cells were observed and images acquired using the Floid^TM^ cell imaging station (Thermo-Fisher) microscope. Images in Figure 2C have been re-coloured for contrast by using the “blue, accent colour 1 light” feature in Microsoft PowerPoint.

#### TK/GCV dose response assay

To determine if TK-expressing virus responds to varying concentrations of GCV to produce a gradient of CPE, Vero E6 cells (10^4^) were seeded in 96-well plates for overnight incubation. Cells were infected with 17D-TK virus at MOI of 0.02 and incubated at 37 °C, 5% CO_2_. Two days after infection, GCV in the concentration range 1 – 25 µM were added with each concentration in n=6 replicates to infected and uninfected control cells. Cells were then incubated at 37°C, 5% CO_2_ and MTS assay performed 5 days post GCV application (7 dpi). Two independent experiments were performed, and each condition assayed in 6 replicates.

#### Synthesis of dF-EdU

dF-EdU is a thymidine analogue described to be preferentially phosphorylated by herpes virus TK over mammalian TK. A detailed description on synthesis and properties of compound dF-EdU has been described elsewhere (Neef et al., 2015). Briefly, the chemotherapeutic drug, gemcitabine (dFdC) is deaminated to form 2-deoxy-2’,2’-difluorouridine (dFdU). The DNA-compatible bioorthogonal functional group ethynyl is attached at the 5’-position of dFdU to yield dF-EDU (Neef et al., 2015).

#### Immunofluorescence assay

Vero E6 cells were seeded at 10^4^ cells per well in 8-well microchamber slides (Ibidi, Germany) for overnight incubation at 37 °C, 5% CO_2_. Cells were washed with PBS and infected with 17D-TK or parental 17D at MOI of 5 and incubated for 1 h at room temperature. Uninfected cells served as negative controls. Cells were then washed with PBS and cultured in DMEM medium (Gibco, Belgium) supplemented with 2% FCS for 2 days at 37 °C, 5% CO_2_. EdU (10 µM) was added to cells and incubated for 2 h at 37 °C, 5% CO_2_. Cells were washed with 3% BSA in PBS and stained following the Click-iT^®^ EdU labelling and detection kit protocol as outlined by manufacturer (Life Technologies, Belgium), Briefly, cells were fixed and permeabilized with 3.7% formaldehyde and 0.5% Triton-X-100 in PBS. Cells were washed twice with 3% BSA in PBS and incubated with the detection cocktail which contains Alexa-Flour 594 labelled copper (Cu) azide, required for the cycloaddition step characteristic of Click-iT chemistry. Cells were incubated with anti-pan flaviviral E protein antibody (4G2) and stained with Alexa-Flour 488 conjugated anti-mouse secondary antibody. Cells were washed with PBS and counterstained with DAPI for nuclear material. Fluorescence images were acquired using Leica SP8 confocal microscope. For dF-EdU Click-iT chemistry assays, similar workflow was followed except for replacing EdU with 50 µM dF-EdU and hamster polyclonal antiserum to yellow fever virus (anti-YF) was used in place of 4G2.

#### Combined TK and GCV animal experiments

Pups with their mothers were put into one of six groups; sham with or without GCV, 17D with or without GCV or 17D-TK with or without GCV. Pups were either sham infected (DMEM media) or with 100 PFU of 17D or 500 PFU of 17D-TK intracranially on day 0. On day 0 and throughout the experiments, mothers of the three groups with GCV were administered once daily with 100 mg/kg GCV (Cymevene) intraperitoneally. Pups of GCV-administered mothers receive a steady dose of GCV through breastfeeding, as described elsewhere (Alcorn & McNamara, 2002). Pups were monitored daily for weight evolution, morbidity, and mortality. Pups were sacrificed seven days after infection, and brains were harvested into RNALater and stored at −80 °C until virus quantification.

#### RT-qPCR

Brain tissues were thawed, homogenized by bead disruption (Precellys®) in Trizol and total RNA was extracted using Zymo Direct-zol RNA extraction kits (Zymo Research) following manufacturer’s instructions. Quantification of YF virus was performed by probe-based RT-qPCR using the iTaq universal probes OneStep kit (BioRad) in a Roche LightCycler 480 PCR machine. YF viral load was calculated using the 2^-ΔΔCq^ method and normalized to β-actin. Primers used are listed in Table S1.

#### Quantification and statistical analysis

Dose response graphs were analysed in Microsoft Excel and graph plotted in GraphPad Prism (version 8) presenting data as Mean+SEM. Differences in weight of pups were analyzed in GraphPad Prism (version 8) calculating Area Under the Curve (AUC) with Holm-Sidak test. Differences with a *p*-value < 0.05 were considered significant.

---

**FIGURE S1 (related to main Figure 2C).**
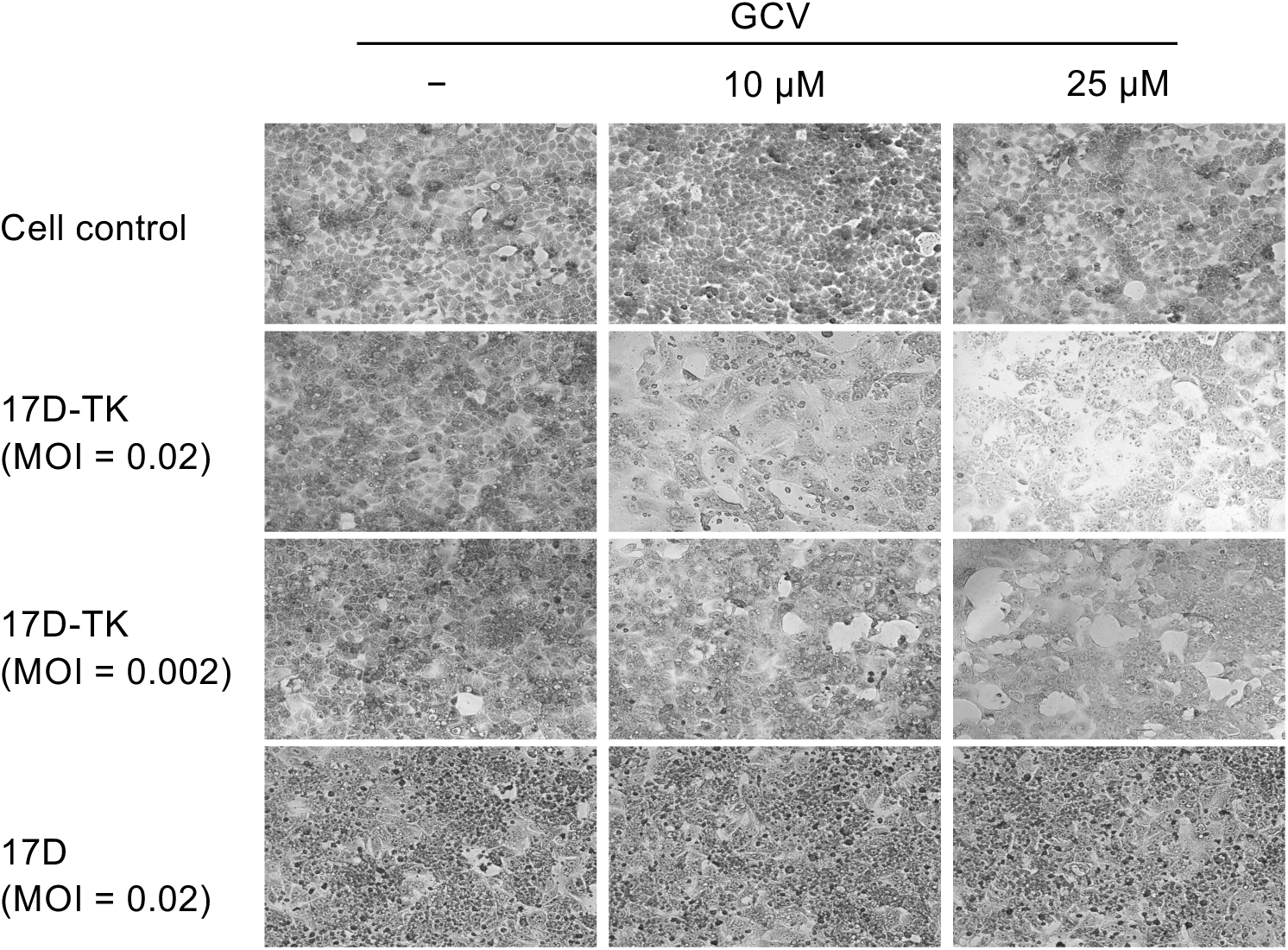
TK expression is required for GCV-induced cytopathic effect. Vero E6 cells were infected with 17D-TK (MOI = 0.02; 0.002) or parental 17D (MOI = **0.02**), and GCV applied 2 dpi at 10 μM or 25 μM. Seven days post infection cells were fixed and stained with methylene blue. Images were acquired using phase contrast microscopy. Representative of two independent experiments. TK, thymidine kinase; GCV, ganciclovir.

**FIGURE S2 (related to main Figure 2C and D).**
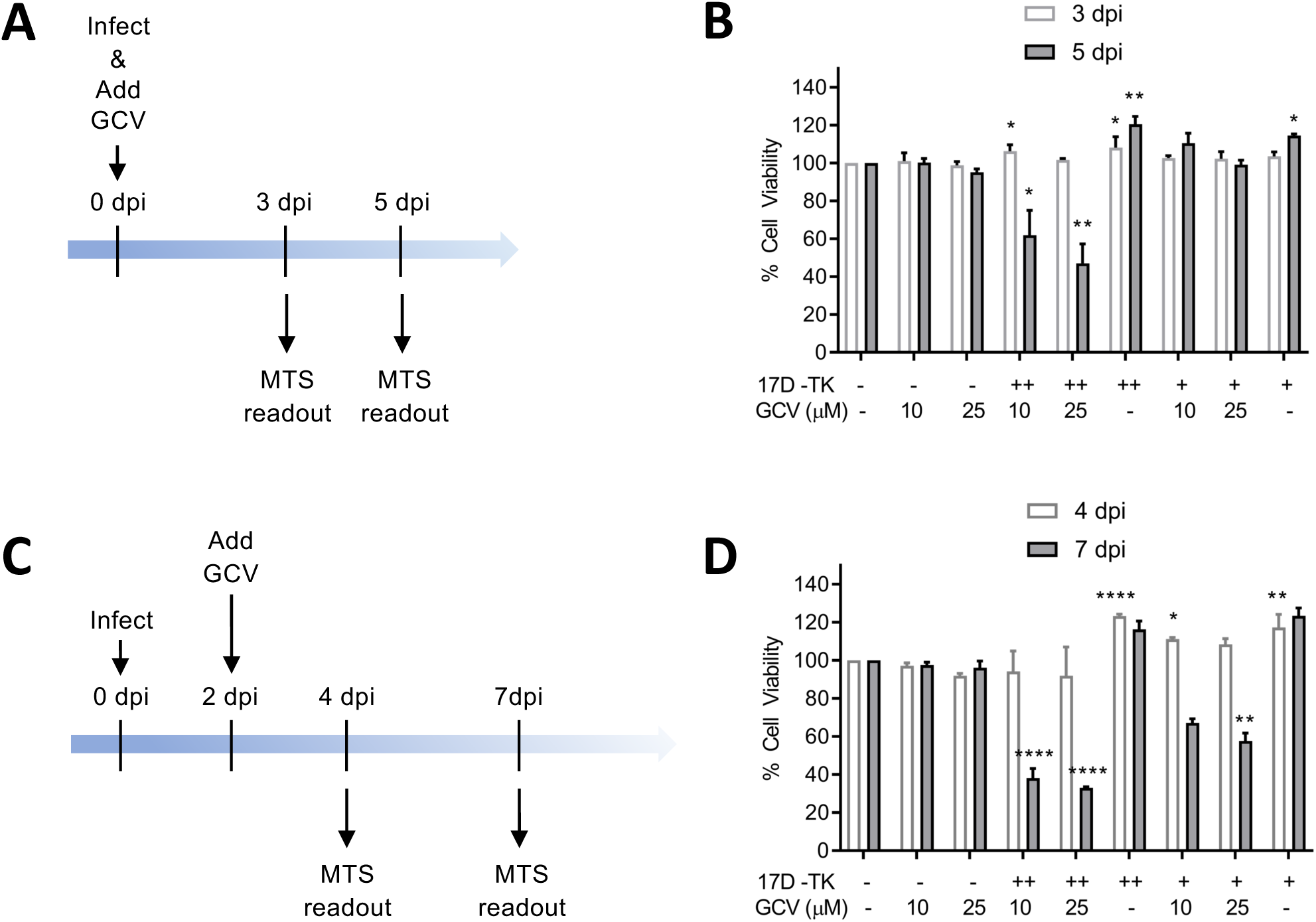
MOI dependence (i.e. proportion of intially 17D-TK infected cells) and time dependence of GCV-induced CPE (i.e. 17D-TK replication). **(A and C)**, schematic depicting infection and assay conditions as in B and D. Vero E6 cells were infected with 17D-TK at a MOI of 0.02 or 0.002 and incubated with 10 μM or 25 μM of GCV, starting on the same day **(A and B)** or 2 days post infection (dpi) **(C and D)**. Cell viability assessed by MTS assay **(B)** 3 and 5 days, or **(D)** 4 and 7 days post infection/GCV application. Uninfected cells without GCV served as controls. Two independent experiments were performed with each experiment performed in n = 6 replicates. Data presented as Mean + SEM. * p<0.05; ** p<0.01; ++, MOI 0.02; +, MOI 0.002.

**Figure S3.**
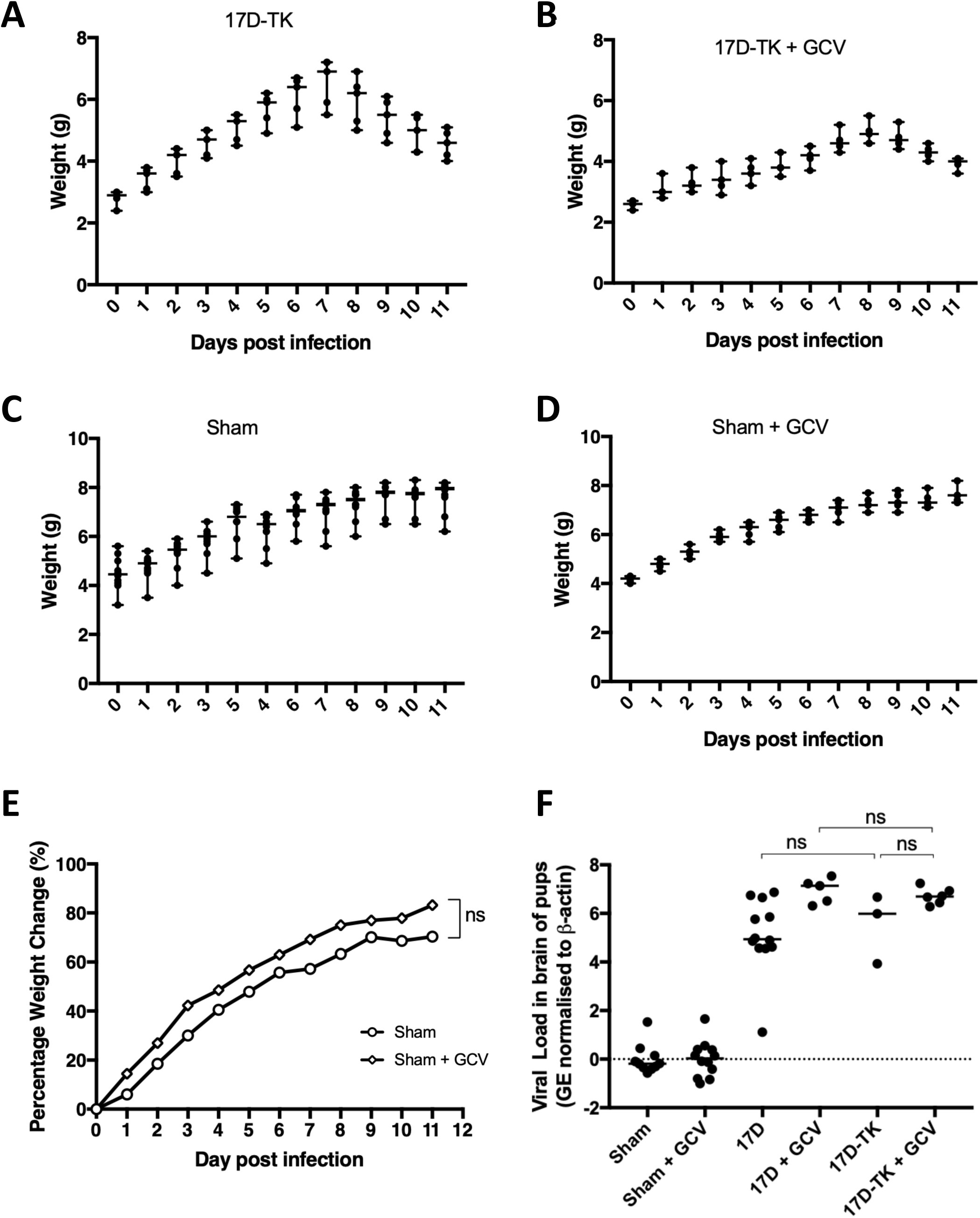
Weight evolution and viral loads in 17D-TK infected pups. (related to main Figure 3). Groups of 4 – 5 days old Balb/c pups were infected with 500 PFU of 17D-TK **(A and B),** or sham infected **(C and D).** Mother of pups in the GCV treatment group [sham + GCV group (n=5); 17D-TK + GCV (n=5)] were daily dosed intraperitoneally with 100 mg/kg GCV while mother of pups in untreated groups [Sham group (n=10); 17D-TK group (n=5)] did not receive GCV. Average weights plotted for sham groups **(E)** to aid direct comparison. **(F)** Viral RNA loads in brains by RT-qPCR.

**FIGURE S4.**
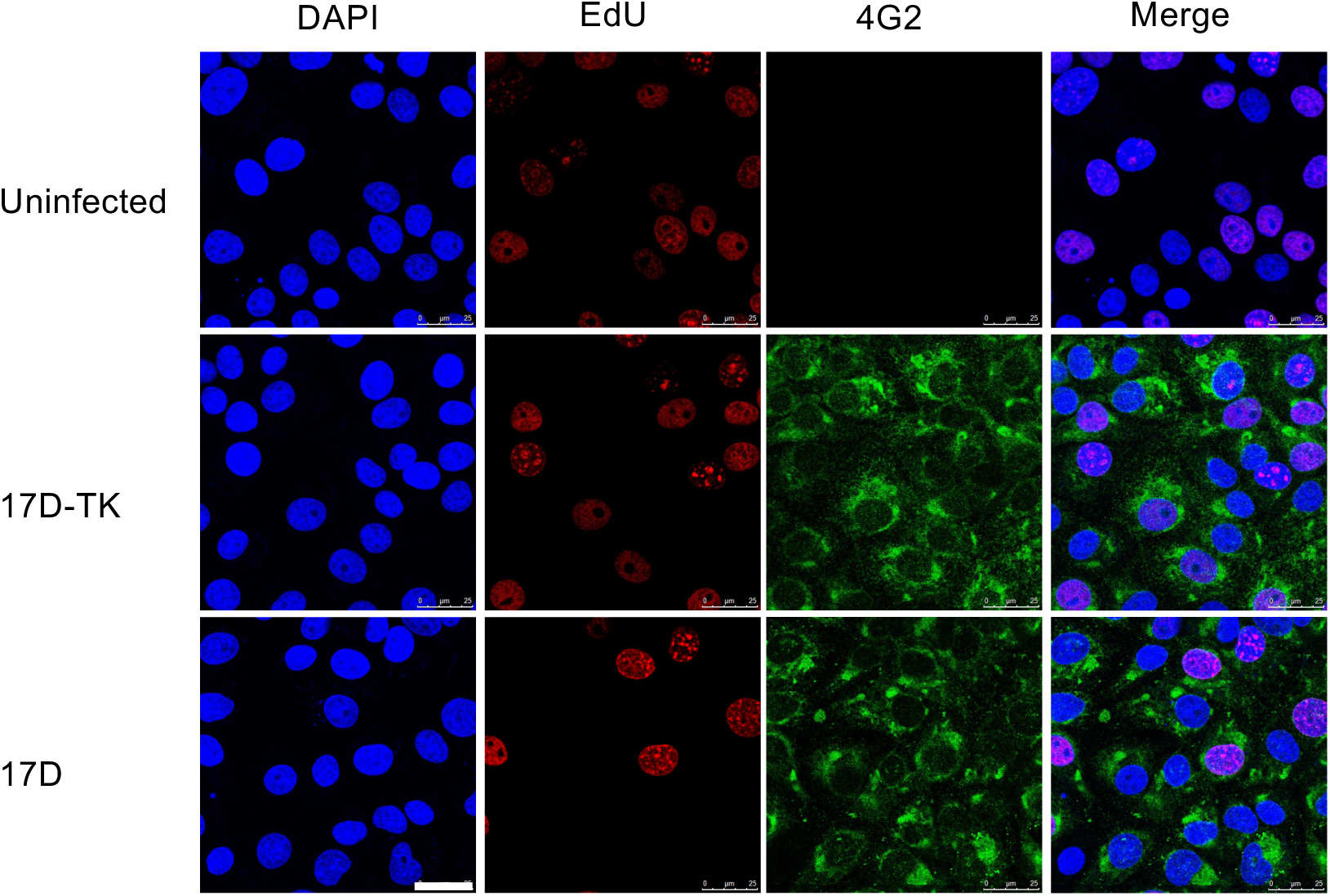
EdU is non-specifically incorporated by cells in synthesis phase independent of virus encoded TK. Vero E6 cells were infected with 17D-TK or parental 17D at a MOI of 5, two days post infection, cells were incubated with 10 μM EdU for 2 hours and stained by Click chemistry using Alexa Fluor-594 conjugated azide reagent. Cells were then incubated with pan-flaviviral anti-Envelope antibody (4G2) and stained with Alexa Fluor-488 conjugated anti-mouse secondary antibody. Images were acquired by confocal microscopy and representative of two independent experiments. EdU; 5-ethynyl-2’-deoxyuridine, white scale bar = 50 μm

**FIGURE S5.**
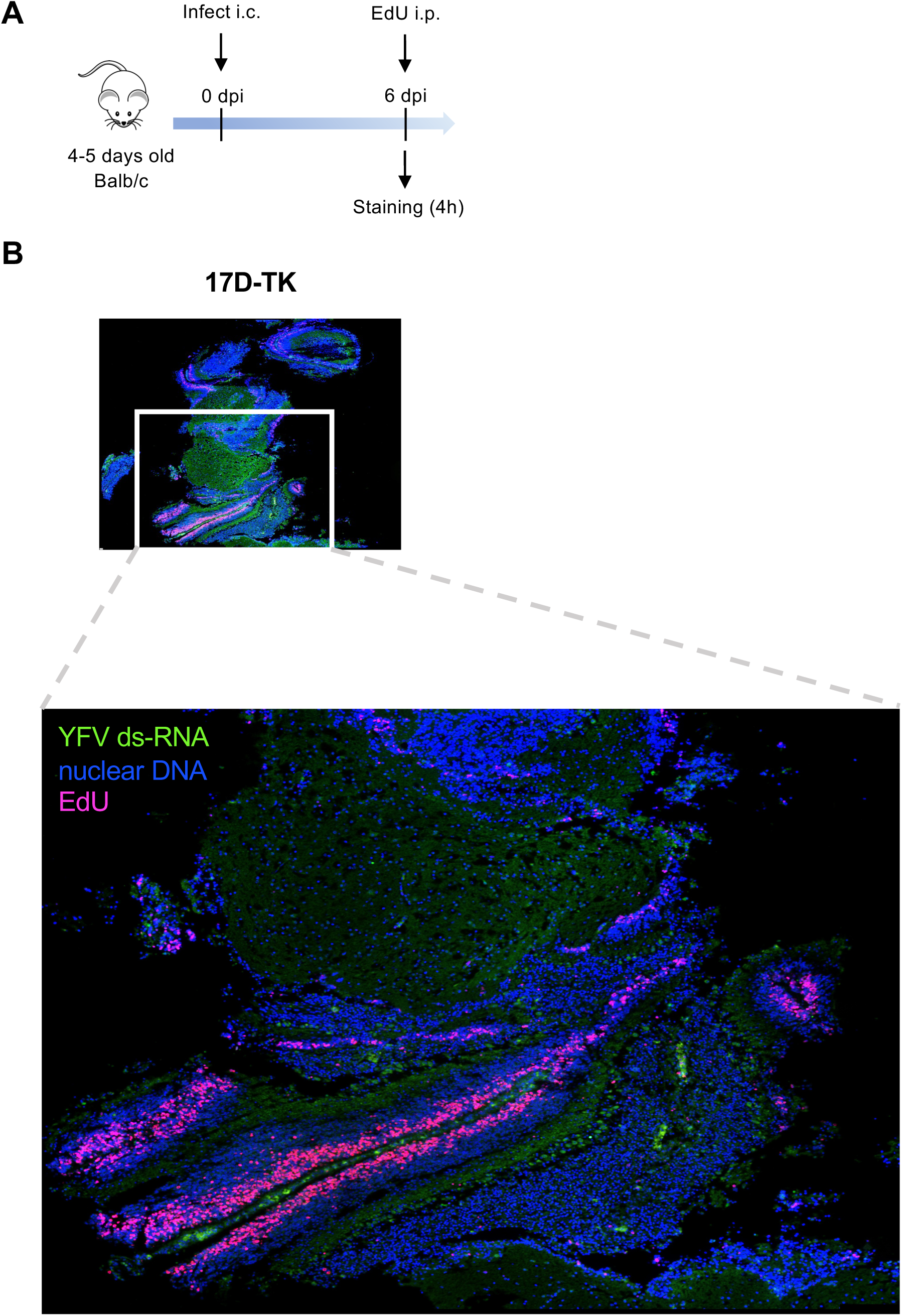
EdU labelling of section from 17D-TK infected brain tissues. (A) Infection and labelling schedule. Balb/c pups were either infected with 500 PFU of 17D-TK intracranially. 6 dpi, pups were injected intraperitoneally with 70 mg/kg EdU and sacrificed 4 hours later. (B) Brains were retrieved, sectioned and stained by Click chemistry. Nuclear DNA is stained with DAPI, YFV dsRNA indicative for viral replication is stained with Alexa Fluor 488 and EdU is stained with Alexa Fluor 594. Images obtained on Leica DMi8 inverted microscope.

