## Supplementary material for "Thymidine kinase-expressing yellow fever 17D reporter virus facilitates prodrug activation and bioorthogonal labelling of infected cells": Key Resources

| Reagent / Resource | Source | Identifier |
|---|---|---|
| **Antibodies** | | |
| Anti-flaviviral E protein antibody (4G2) | Sigma Aldrich | MAB10216 |
| Alexa-Fluor 488 anti-mouse secondary antibody | Invitrogen | A-11001 |
| Polyclonal mouse anti-yellow fever virus antiserum | Kai Dallmeier, KU Leuven | N/A |
| Anti-dsRNA monoclonal antibody J2 | Scicons | RNT-SCI-10010200 |
| DAPI | Sigma Aldrich | D9542 |
| **Plasmids** | | |
| BAC pShuttle-17D | Kai Dallmeier, KU Leuven | N/A |
| **Viral strains** | | |
| Yellow Fever 17D Virus (17D) | Recombinant virus derived from pShuttle-17D infectious clone | |
| Yellow fever 17D virus-TK reporter (17D-TK) | This paper | |
| **Chemicals** | | |
| Dolethal | Vetoquinol, SA | |
| Agarose | Invitrogen, USA | |
| Paraformaldehyde | Sigma Aldrich | #158127 |
| Phenazine methosulphate (MTS reagent) | Promega | G1111 |
| Ganciclovir (Cymevene) | | |
| dF-EdU | Nathan Lüedtke; McGill University (Neef et al., 2015) | N/A |
| **Critical Commercial Assays** | | |
| TransIT-LTI transfection reagent | Mirus Bio | MIR 2300 |
| Click-iT EdU labelling and detection kit | Life Technologies, Belgium | C10337 |
| Direct-zol RNA extraction kit | Zymo Research | R2052 |
| iTaq universal probes one step PCR kit | Biorad | #1725141 |
| **Cell Lines & Animals** | | |
| Hamster: BHK-21J cells | Peter Bredenbeek, LUMC, NL | N/A |
| African Green monkey: Vero E6 cells | Peter Bredenbeek, LUMC, NL | N/A |
| Balb/c mice | Janvier Labs, France | N/A |
| **Media** | | |
| DMEM | Gibco, Belgium | 41965-039 |

| Reagent / Resource | Source | Identifier |
|---|---|---|
| MEM-2X | Gibco, Belgium | 21935-028 |
| FCS | Hyclone | SV30160 |
| 2mM Glutamine | Gibco, Belgium | 25030-024 |
| **Oligonucleotides** | | |
| YFV-NS3-Forward-P2: | CACGGCATGGTTCCTTCCA | |
| YFV-NS3-Reverse-P2: | ACTCTTTCCAGCCTTACGCAAA | |
| YFV Probe-P2 | MFAM-CAGAGCTGCAAATGTC | |
| EHV4-TK-F (Primer #886) | TGTCTGGTCGTAAAGCTCA | |
| EHV4-TK-R (Primer #1369) | GTGTCTTACCCTGGGCTTTGCG<br>GCCGCTAGGACCGGGGTTCTC<br>CTCCACGTCGCCACAGG | |
| b-actin-F | GGCATCCACGAAACTACCTT | |
| b-actin-R | AGCACTGTGTTGGCGTACAG | |
| **Software & Algorithms** | | |
| Graphpad Prism version 8 | Graphpad software | www.graphpad.com |
| Floid cell imaging station | Thermo-Fisher | #4471136 |
| Confocal microscope, SP8 | Leica | |
